# Evidence against a cortical module for processing communicative gaze

**DOI:** 10.1101/2023.01.20.524674

**Authors:** Marius Görner, Peter W Dicke, Peter Thier

## Abstract

Where we look lets others know what we are interested in and allows them to join our focus of attention. In several studies our group investigated the neuronal basis of gaze following behavior in primates and described a *gaze following patch* (GFP) as the underlying functional unit. This makes the GFP a promising neurobiological correlate of Simon Baron-Cohen’s *eye direction detector* (EDD), an integral part of his *mindreading system*. With the latter, Baron-Cohen proposed a set of domain-specific neurocognitive modules implementing a Theory of Mind - the attribution of mental states to others. The tenet of domain-specificity requires that the EDD processes exclusively eye-like stimuli. In this preregistered fMRI study, we aimed to critically test if the GFP fulfills this criterion. Contrary to previous studies, we found that this was not the case and that it functionally and anatomically overlapped with the MT+ complex, putting the plausibility of Baron-Cohen’s *mindreading model* as a set of domain-specific modules into question.

**Teaser:** The presented results demonstrate that a brain area previously thought to be a domain-specific cortical module specialized in processing other people’s focus of visual attention is, in fact, domain-general. More specifically, we show that it is indistinguishable from cortical areas involved in the processing of visual motion. These findings add to a growing body of evidence arguing against domain-specific accounts in social neuroscience which posit that the functional building blocks of social behaviors are facilitated by dedicated neurocognitive modules.

## Introduction

The direction of our gaze plays a fundamental role when we interact with a fellow human. Just by observing the other person’s gaze averting her eyes from one towards a new location, we can tell that her attention has been grabbed by something else, inevitably also attracting our own attention. Thereby we establish *joint attention-* the congruence of attentional foci. Further, by mapping our own beliefs, desires and intentions associated with the object of the otheŕs attention enables us to develop a *theory of (the otheŕs) mind* (ToM) which allows us to predict the otheŕs behavior. With his *mindreading system*, Simon Baron-Cohen suggested a functional framework which is implemented by four domain-specific neuro-cognitive modules underpinning the route towards a full-fledged TOM. In this, the *eye direction detector* (EDD) is the node that contributes the information on gaze direction which is integrated with information from an *intentionality detector* (ID) and relayed to the *shared attention mechanism* (SAM), a term often used as an alternative to *joint attention* in order to emphasize the agent’s awareness of the congruence of their attention foci. Finally, the latter passes its information to the *theory of mind mechanism* (ToMM)^1–3^. He based this idea on Fodor’s concept of modularity, which implies that the modular components have distinct and specific neural architectures. These architectures would consist of populations of neurons that exclusively process the respective information; in the case of the EDD this means eye-like stimuli with their typical shape and motion properties^1,4^. Apart from Baron-Cohen’s concept, the idea of modularity is widespread in the field of social cognitive neuroscience^5–8^ but has also been criticized from early on^9–11^.

Given the ecological importance of judging gaze directions – whether for predator avoidance, prey detection or other forms of social communication^12^ – it seems indeed conceivable that a specialized neural mechanism evolved to optimize processing in order to facilitate rapid behavioral responses to the otheŕs gaze (see^13^ for a recent review on the evolutionary perspective on gaze-following). How fast you can detect a staring predator may decide upon life and death. Support for this idea is provided by behavioral studies showing that responses to gaze stimuli in both humans and macaques appear to be reflex-like, i.e. they are fast and cannot be suppressed^14–16,6,17–19^. Perrett and colleagues were the first to describe neurons in the superior temporal sulcus (STS) that are tuned to varying gaze, head and body orientations^20^, yet without testing these neurons for behavioral relevance. Other studies have reported related results on the representation of gaze stimuli in the temporal cortex of monkeys and humans, in the latter emphasizing the right temporal cortex^21–23^, which in general dominates the processing of information provided by faces^24^ (see^25–27^ for reviews).

In a series of behavioral, fMRI, and electrophysiological studies our group investigated the neural basis of gaze-following behavior in macaques and humans. In this work we were able to identify a circumscribed region in the posterior temporal cortex of humans and macaques, which we baptized the *gaze following patch* (*GFP*). This region is comparatively more active during gaze-following than in control conditions, characterized by very similar visual and motor demands, yet lacking the need to follow gaze^19,28–34,8^. This characteristic makes the GFP an obvious neural correlate of Baron-Cohen’s EDD. The distinctive feature of our experimental design was that it required *active* gaze-following, i.e. participants did not only passively observe a face gazing in a certain direction as in the previous work mentioned but had to utilize it to identify a target among distractors and to subsequently shift their own visual attention to it. In the control condition that we used the target had to be identified based on the iris-color^31^ or the identity^8^ of the stimulus face. We found that the GFP –– revealed by contrasting gaze-following and iris-color matching –– does not overlap with any of the neighboring elements of the face patch system^5,31,35^, albeit positioned close to the middle face patch. Moreover, it is located close to the middle temporal area (MT), well-known to be instrumental in the processing of visual motion. In fact the human GFP is overlapping with parts of the MT+ complex delineated by the neurosynth.org database^36^ as well as by our own pilot experiments involving low-level motion stimuli in humans (see Figure 2 and S1). Visual motion may indeed have been a critical stimulus component in our work as the gaze stimuli used were based on a portrait in which the eye axes switched from straight ahead to averted, thereby giving the impression of a gaze saccade – i.e., high speed motion of the eyes. Although the control condition involved the same motion component, it was behaviorally irrelevant as it was the iris-color that participants had to exploit to identify the target with the matching color. Thus, it may well be that the critical aspect of the stimulus that triggered the GFP activity was not the actual retrieval and use of gaze direction but the need to attend to the associated movement of the eyes, activating generic motion specific cortex.

Hence, in the present, pre-registered study^37^, we tested whether the GFP needs to be fused with the visual motion processing system, thereby depriving the GFP of its property of domain-specificity assumed by the earlier work^8,28,29,31,38^. This seems to be the case: our results clearly show that not only the observation of an eye saccade is accompanied by activity in the GFP. Rather, a stimulus that bears no visual resemblance to eyes, but has a movement pattern comparable to a saccade, also elicits activity. Furthermore, we show that the activity in the GFP cannot be distinguished from activity in a ROI delineated by low-level visual motion stimuli, as already suggested by the anatomical congruence (see Figure 2). Given that all of our experiments have been designed to reveal an EDD as an example of a cortical module underpinning domain specificity, we propose that our results have strong implications beyond the ontological status of the GFP. Considering that the GFP ––the best candidate of a cortical realization of the EDD so far––failed a critical test of domain specificity suggests that gaze-following may not have a domain-specific neural basis at all, but instead be grounded in domain general cortical processing serving a much wider range of perceptual functions.

## Results

### Confirmatory Experiment 1

In each trial of the first experiment the participants got to see a gaze shift as well as a change of the iris color of the demonstrator face, and a rotation of cubes that were positioned on the face. Auditory instructions indicated which of the cues should be used to identify the target (see Figure 1). For each participant we used the BOLD contrast between trials in which they had to use the gaze shift and trials in which they had to use the change of the iris color to identify the GFP-ROI. For all participants except sub-06 we could identify individual GFP-ROIs, for sub-06 we used a spherical ROI that was centered on the local maximum closest to the reference coordinates of the GFP (Figure 3/ S1 for a depiction of the ROIs). Using these as masks, we extracted the beta values of the *gaze-shift* and *cubes-rotation* conditions for each run of each participant (Figure 3).

**Figure 1.**
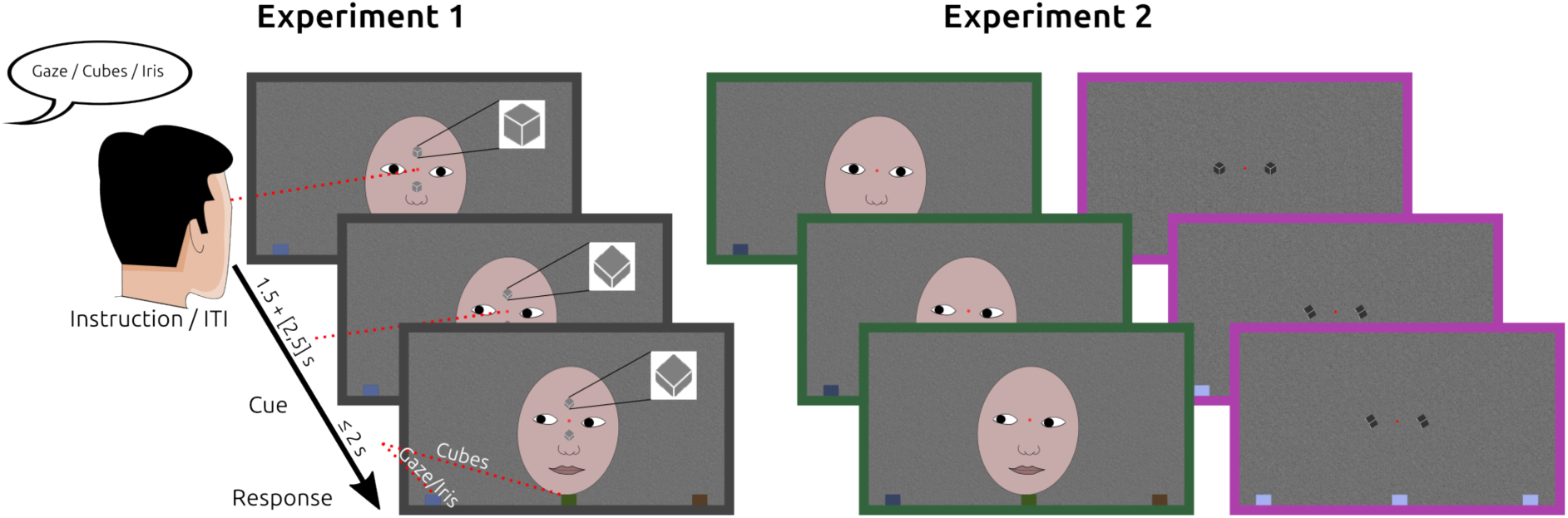
Illustration of experiment 1 and 2. In the example of experiment 1 the cubes indicate the middle target and the gaze as well as the iris-color the left target. Depending on the instruction, the participant will have to choose the respective target. In experiment 2 the two stimulus components gaze-shift and cube-rotation are presented separately. In the experiment the schematic face shown here is replaced by a photo of a person.

**Figure 2.**
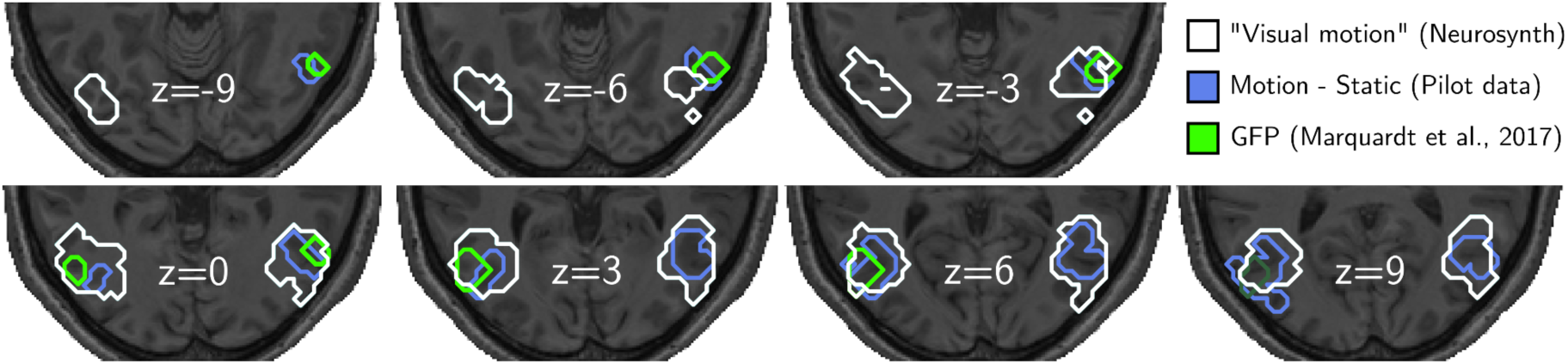
Anatomical locations of motion related areas and the GFP. Shown are transversal cuts from *z* = – 9 to *z* = 9 in steps of 3 *mm*. Locations given by the neurosynth^36^ database when searching for “visual motion” are outlined in white. The result of the MT localizer of Pilot 1 is outlined in cyan and the GFP from Marquardt et al ^31^ is outlined in green.

**Figure 3.**
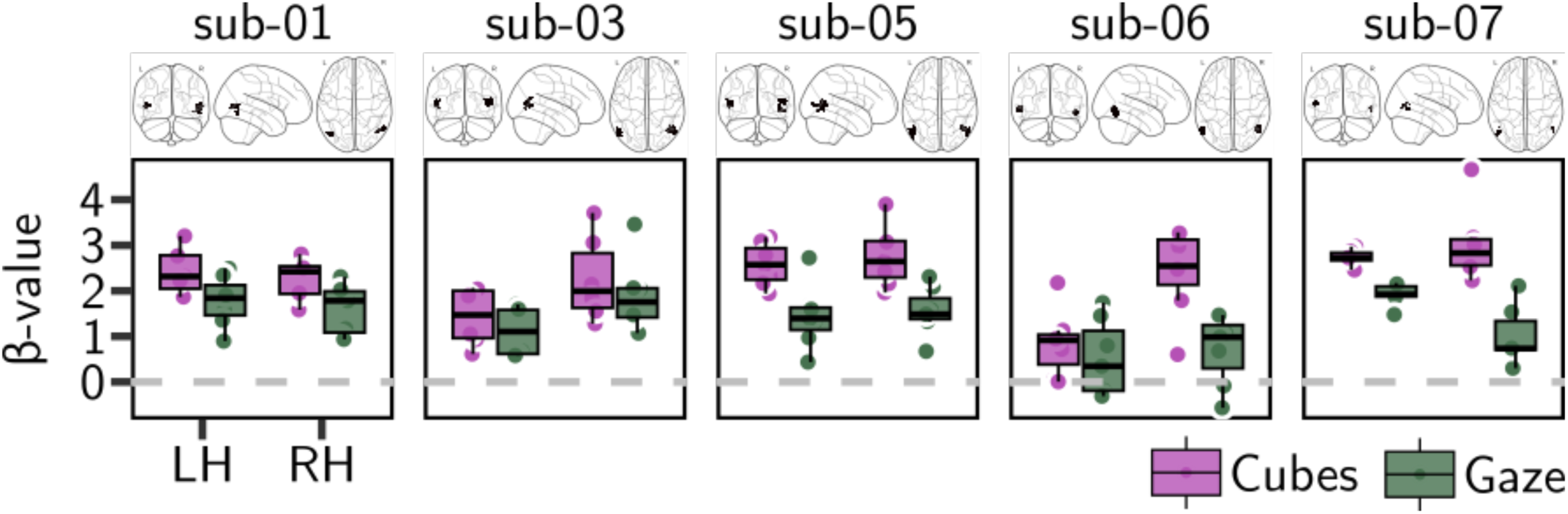
For each participant beta values extracted from the GFP-ROIs are shown grouped by hemisphere and condition. Individual ROIs are shown on top of each boxplot For all but participant 6, the GFP could be identified individually. In both conditions the GFP is consistently activated with beta values larger than 0. Beta values related to the *cube-rotation* are consistently larger than those related to the *gaze-shift* condition.

Statistical analysis supports a difference in the activity associated with the two conditions, but with larger activity in the *cubes-rotation* condition than in the *gaze-shift* condition (see Table 1, mixed effects model with *run* as group-level factor, *BFs* are *log_10_* transformed).

**Table 1.** Statistical results regarding differences between conditions (Table S1, Q1).

| | $\Delta\beta^{\text{Gaze-Cube}}$ | $CI^{95\%}$ | $BF_{01}^{Q1}$ |
| --- | --- | --- | --- |
| LH | -0.65 | $[-0.87, -0.4]$ | -11.02 |
| RH | -1.15 | $[-1.48, -0.82]$ | -88.93 |

The non-logarithmically transformed Bayes Factor for the right hemisphere exceeds numerical precision, fluctuating between positive and negative values, meaning that the posterior probability of 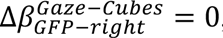, i.e., no difference between the BOLD responses in the conditions.

Testing for differences between the left and the right GFP within each condition the data support the existence of a difference for the *cube-rotation* with *moderate* evidence but equal activation related to *gaze-shift* with *strong* evidence (see Table 2, mixed effects model with *run* as group-level factor, *BFs* are log transformed).

**Table 2.** Statistical results regarding differences between hemispheres (Table S1, Q2).

| | $\Delta\beta^{RH-LH}$ | $CI^{95\%}$ | $BF_{01}^{Q2}$ |
| --- | --- | --- | --- |
| **Gaze** | 0.01 | [-0.28, 0.31] | 1.41 |
| **Cubes** | 0.54 | [0.21, 0.88] | -0.65 |

### Experiment 2

Unlike in the first experiment, in this second experiment participants saw only either the cubes or the demonstrator face shifting eye-gaze in a given trial. Using the individual GFP-ROIs from experiment 1 and the individual MT+ ROIs from the motion localizer tasks (Figure S1 shows all individual ROIs) we estimated hemodynamic response functions (*HRF*) for each participant individually as well as the group level *HRFs* grouped by ROI, hemisphere, condition and target (left, middle, right) (see Figure 7).

**Table 3.**
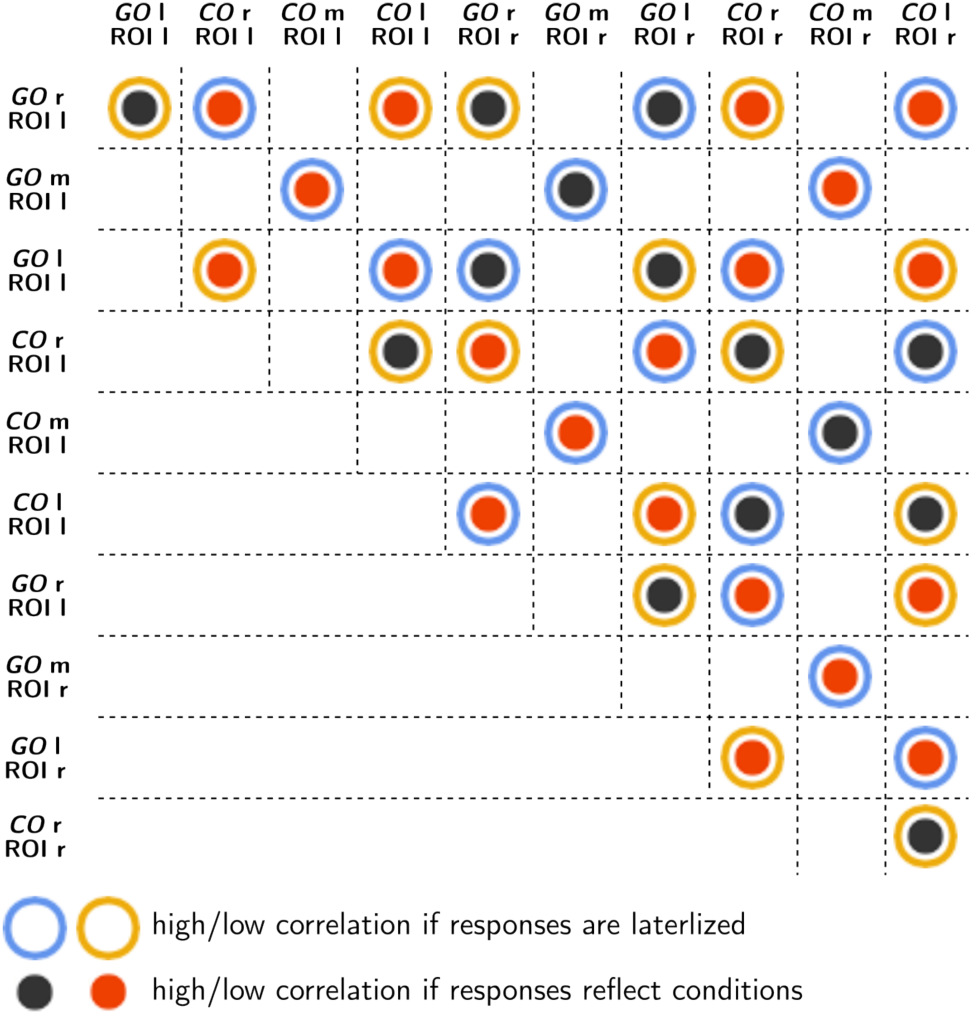
Predictions of magnitudes of correlations. GO = gaze-only, CO = Cubes-only; l – left target, m – middle target and r – right target. Correlations coefficients will be computed for all marked pairs. For pairs marked by blue circles correlation coefficients are expected to be higher than for those marked by yellow circles if responses are lateralized and triggered by low-level motion components (Q3.1/ 4.1). If, however, responses reflect the conditions (Q3.2/ 4.2), then correlation coefficients for pairs marked by dark gray dots will be greater than those marked by red dots. Target-2 trials (middle) are only paired with itself to consistently maintain lateralization with respect to the left and right targets. The analysis will be carried out on data of the GFP and MT+ (ROI = [GFP, MT]). Left and right refers to the hemisphere.

There were clear responses for all conditions in all ROIs and hemispheres with slightly larger amplitudes in the *gaze-only* related responses. The temporal patterns are indistinguishable when comparing within each ROI across conditions, as well as when comparing within conditions across ROIs. Especially when comparing the responses of one condition across ROIs of the same hemisphere, across columns in Figure 4, there is virtually no difference. One characteristic of the responses that stands out is that in all plots (except for the right GFP) there is one curve that is different from the other two. The difference is in the amplitude as well as the timing of the peak and trough. For the left ROIs it is the response related to the right target and for the right ROIs the one related to the left target. This pattern corresponds to the prediction based on the assumption that the responses in the ROIs are lateralized, which is a property of area MT (Table S1, Q3.1 & 4.1).

**Figure 4.**
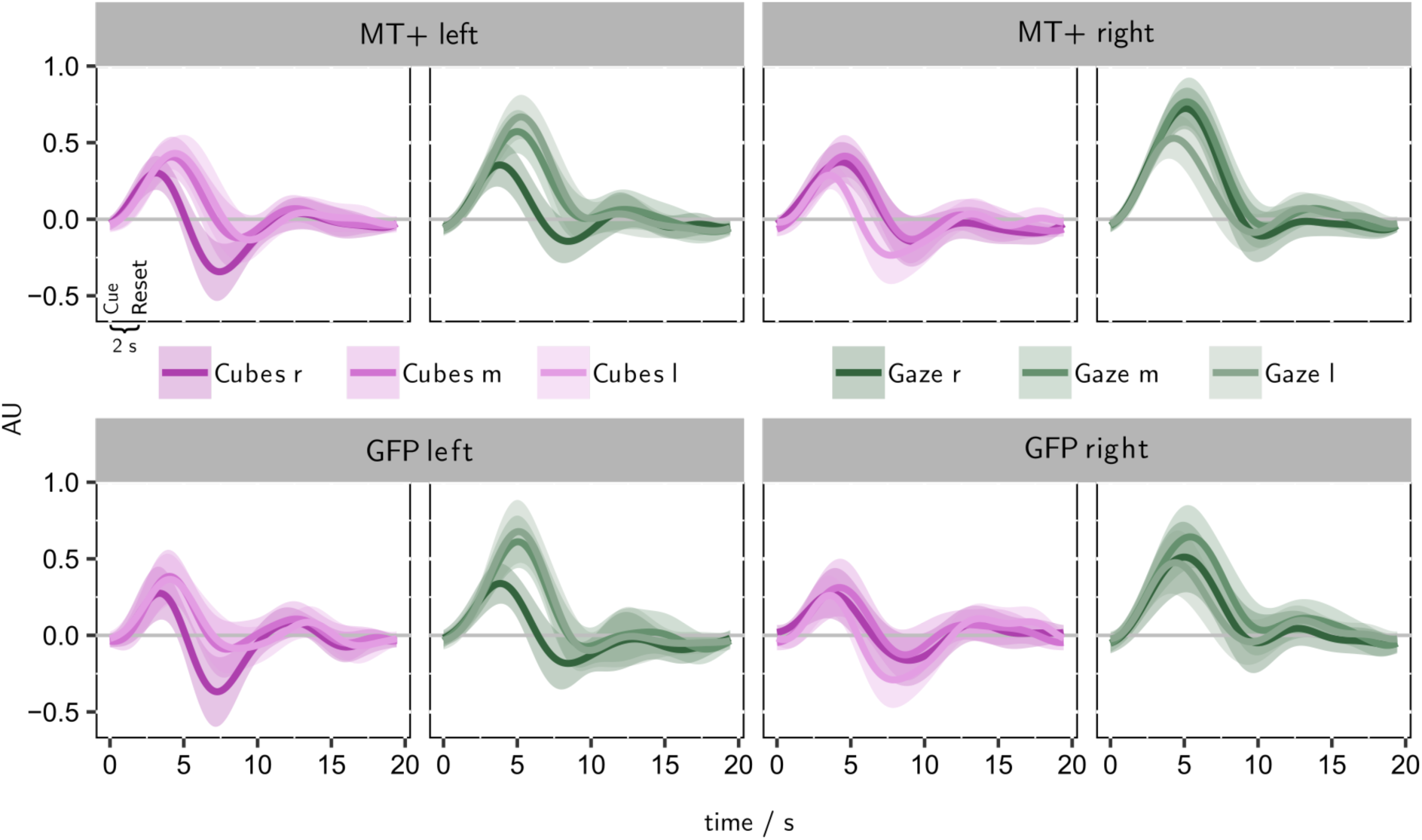
Group-level HRF estimates grouped by ROI, experimental condition and target ID. Especially in both ROIs of the left hemisphere the pattern predicted based on a lateralized BOLD response to visual motion is apparent. In these two ROIs the response to the right target is diminished compared to the other two response curves. In the ROIs of the right hemisphere the effect is less obvious, the pattern, however, is present, as well.

To quantify the similarity between the responses to the *gaze-only* and *cubes-only* conditions we computed the correlation between pairs of responses (Table S1, Q3&4). The correlations were computed for each participant individually. The relevant pairs for which we computed the correlations are shown in Table 3. Correlations between *HRF*s were only computed within each ROI (GFP or MT+), however, across hemispheres and conditions. To directly compare the GFP with MT+ the obtained correlation coefficients were then analyzed for their correlation, i.e., asking the question if the correlation coefficients for the pairs of *HRF*s have similar values in the GFP and MT+ (Table S1, Q5). Figure 5 shows the results of the direct comparisons between the two ROIs (Table S1, Q3-5). Data of the participant for whom we could not identify an individual GFP-ROI are shown in the scatter plot of Figure 5 with reduced opacity. We did not include this participant’s data in the statistical analysis presented here because the fact the ROI could not be inferred directly from the *gaze-shift* minus *iris-color* contrast image casts doubt on whether there was a task-related signal at all. However, including this subject changed the results only quantitatively. Our central finding is that both ROIs showed the same correlation pattern. We found *strong* evidence that both, the GFP and MT+, exhibited differences between the groups reflecting conditions (black and red bar plots in Figure 5; Table S1, Q3.2 & 4.2). Questions 3.1 & 4.1, asking whether the ROIs show lateralized responses independent of the experimental condition, cannot be answered with sufficient evidence (see *Methods - Transparent Changes*); the GFP data, however, suggests with *moderate* evidence that there is no difference between the groups reflecting lateralized responses (*BF*_01_ = 0.6; Figure 5, top left, blue and yellow bars). The similarity of the correlation patterns for the two ROIs is also expressed in their direct comparison. The correlation between the correlation patterns for the GFP and MT+ is 0.75, meaning that if a given pair of HRFs is (not) correlated in the GFP it is also (not) correlated in MT+ (Figure 5, scatter plot, Table S1, Q5).

**Figure 5.**
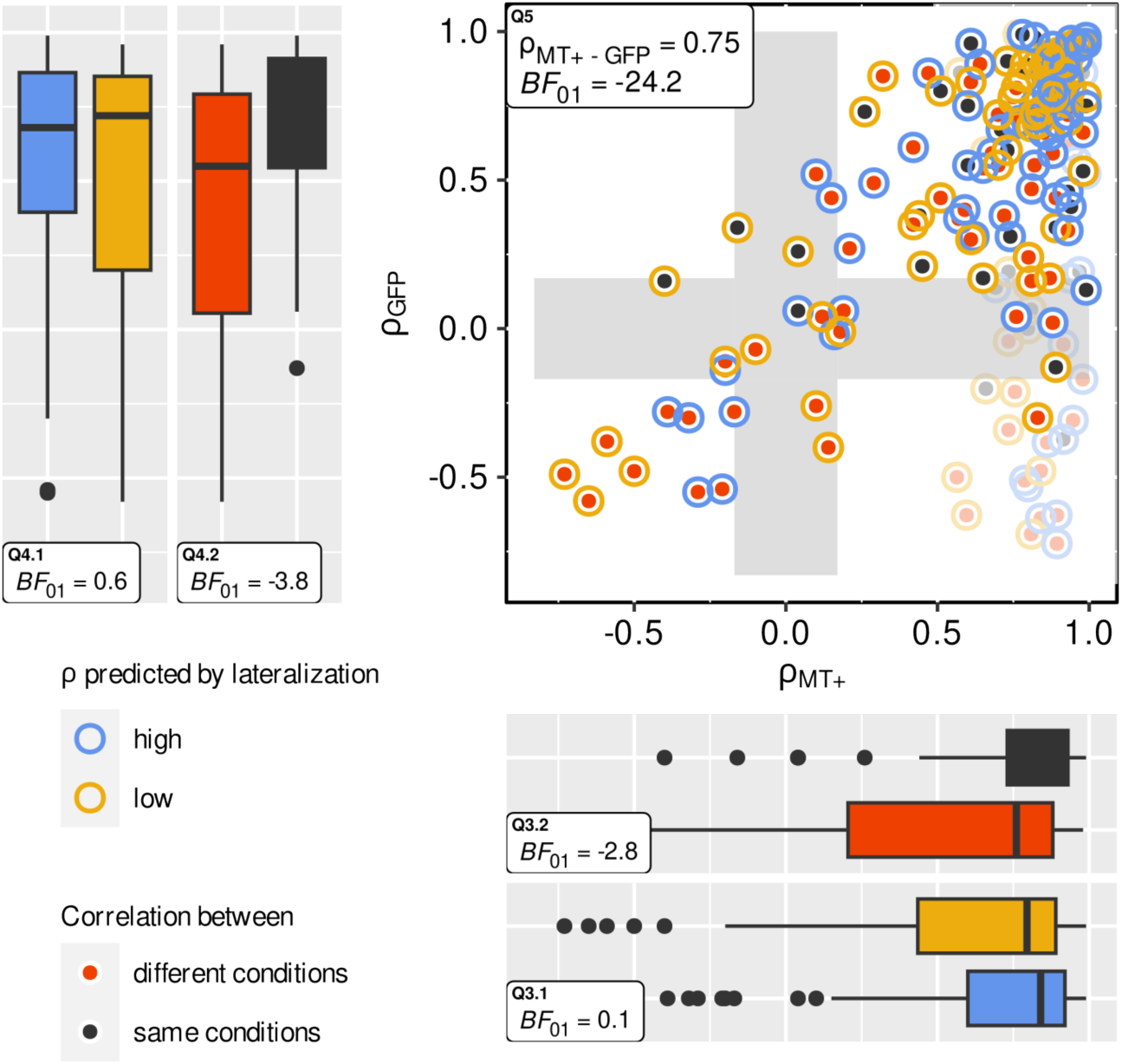
Results of the correlation analysis. The boxplots illustrate the correlation coefficients grouped according to the two predictions (blue-yellow: Q3.1/ 4.1; red-black: Q3.2/ 4.2) for the two ROIs (top-left: GFP; bottom-right: MT+). The inlets show the log_10_ transformed Bayes Factors. The scatter plot shows the correlation coefficients based on the two ROIs independent of the groupings. It can be seen that if a pair of HRFs is correlated in one ROI, it tends to be correlated in the other as well. The inlet shows the correlation coefficient between the two ROIs as well as the respective Bayes Factor indicating *extreme evidence* in favor of a correlation different from 0. The data points plotted with a lower opacity come from the subject for whom it was not possible to identify an individual GFP-ROI and whose data was not included in the quantitative analysis.

### Exploratory

To explore brain wide qualitative differences between the activity related to the *gaze-shift* and the *cube-rotation* conditions we looked for brain areas where one of the experimental conditions elicited positive *β*-values and the respective other condition non-significant (*p* ≥ 0.05*, uncorrected*) or negative *β*-values. Further, we applied a cluster size threshold with a minimum of 15 adjacent voxels to remove noise and focus on broader activity patterns. Areas shaded in green in Figure 6 are those that carried positive *β*-values during *gaze-shift* following and non-significant/ negative *β*-values during *cube-rotation* following. Areas shaded in purple have the opposite pattern. Finally, the uncolored areas are those in which the *β*-values were not significant or both conditions evoked a response of the same sign.

**Figure 6.**
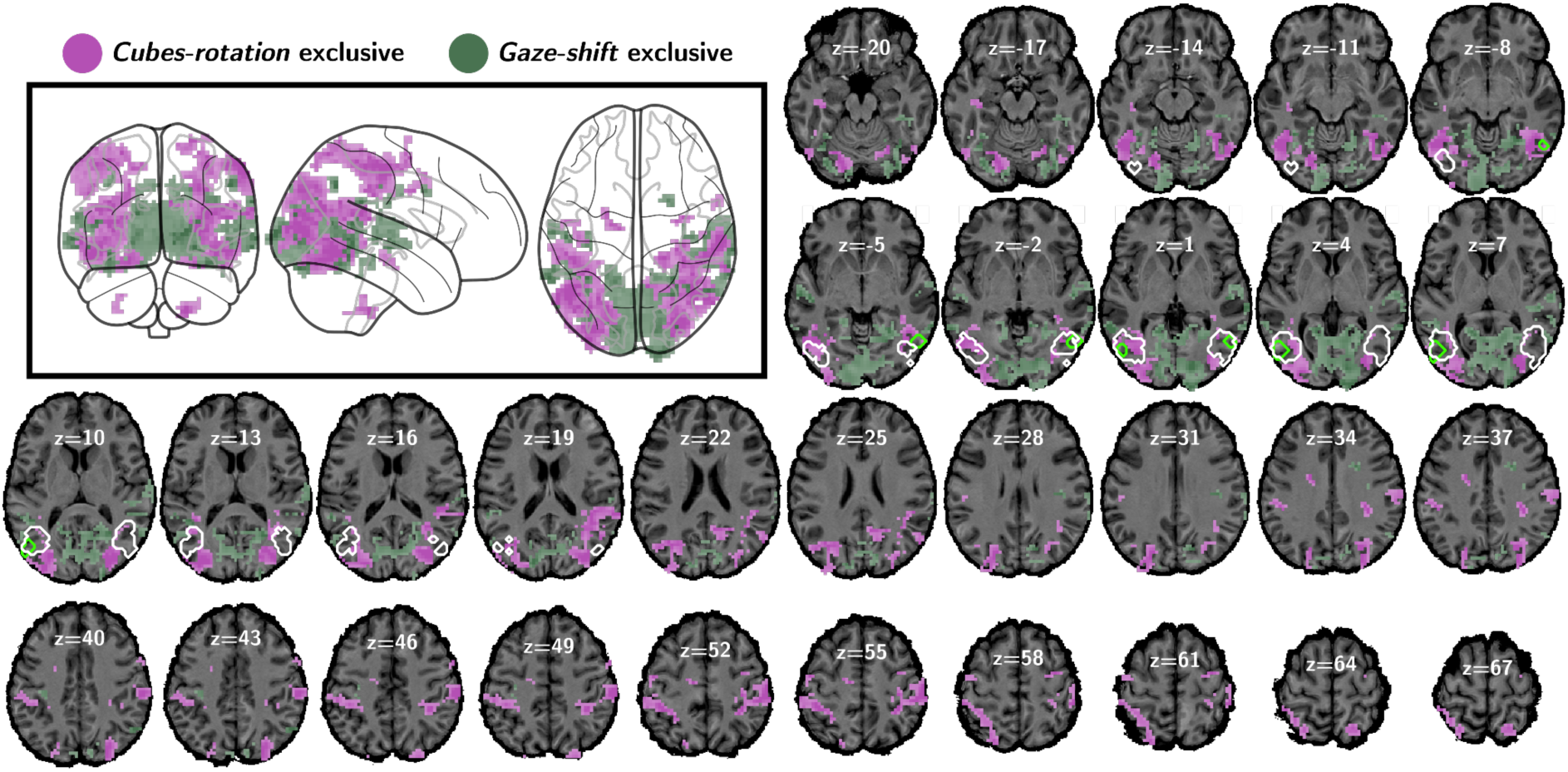
Exploratory whole brain analysis of group-level activity patterns exclusively associated with *gaze-shift* and *cube-rotation* following. First, the *β*-maps were thresholded with p=0.05 (uncorrected) and a minimum cluster-size of 15 voxels. Then all voxels that were positively correlated with one condition were removed from the *β*-map of the other condition. I.e. all areas shaded in green represent activations that were positively correlated only with the *gaze-shift* condition, and all areas shaded in purple were positively correlated only with the *cube-rotation* condition. The area outlined in bright green is the GFP-ROI based on data from Marquardt et al. ^31^. The areas outlined in white the MT+-ROI based on the neurosynth database.

In the context of our experiment, areas exclusively associated with gaze shifts tended to be located in more medial parts of the occipital cortex and in the temporal cortex, whereas areas exclusively associated with cube rotations tended to be located in lateral parts of the occipital cortex and extend into parietal areas. More specifically, hotspots related to gaze shifts were located in the medial parts of early visual areas BA 17, 18 and 19. Furthermore, there were hotspots in BA 37 (more medial) in both hemispheres, in BA 23 and in the temporal cortices with hotspots in BA 41, 22 and 21. Areas exclusively activated in the *cube*-*rotation* condition were located in early visual areas BA 18 and 19, as well as in BA 37 (more lateral). In the parietal cortex, right BA 39 (extending into BA 22) and bihemispheric BA 40 and BA 7 exhibited hotspots.

## Discussion

It has been debated for many years whether there are neurocognitive modules that underlie complex (social) behaviors^39,9–11,40^. The discussion was sparked by Jeremy Fodor’s influential thoughts on the modularity of the mind. But in fact, Fodor distinguished between *input systems* and *central systems* and proposed that only the former were modular in the strict sense^4^. Ignoring this distinction, social cognitive neuroscience has proposed central neurocognitive modules processing different aspects of social interactions. Simon Baron-Cohen’s *mindreading system* is a particularly influential case in point.

In this study, we set out to critically reexamine the assumption of a modular architecture of mind-reading by questioning the existence of an EDD, a putative cortical module underpinning the perception of gaze direction. Such a module has been suggested to be accommodated by a distinct region in the occipitotemporal cortex (the *gaze following patch*, GFP)^1,2,28,31,34,38^. In order to challenge the assumed specificity of the GFP for the otheŕs gaze, we tested its response to a stimulus that carried directional information as well as visual motion while not resembling eyes. As a first step, we reproduced the GFP in each experimental subject by computing the functional contrast *gaze-shift following* minus *iris-color* that had been used in previous studies to delineate it. We then compared the *β*-values associated with the control condition lacking eye features, *cube-rotation following* with those related to *gaze-shift following* for the GFP-ROIs. This comparison clearly showed that the GFP is active not only when observing a gaze shift of a demonstrator, but also when observing a non-biological stimulus that has similarity to eyes, while moving in a comparable way and having a comparable behavioral relevance in the context of the experimental task. In fact, the *β*-values associated with the *cube-rotation* condition were consistently larger than those related to the *gaze-shift* condition for all subjects. This result speaks against the hypothesis that the GFP is a cortical module that processes others’ gaze direction in a domain-specific manner. We can think of two reasons why the activity associated with the *cube-rotation* condition is even greater: either the detection of the cube rotation was more difficult – potentially due to interference with the face/ gaze stimulus in the background –, or the motion energy of the cube rotation was larger. The latter could be due to the fact that the cube rotation was animated by several frames, whereas the gaze shift was animated by only two frames. Whichever option may hold, both would indicate even more strongly that the GFP is not a dedicated EDD, since its activity scales either with the difficulty of detecting visual motion or with the energy of the motion stimulus. The fact that the activations associated with cube rotations were consistently greater than those associated with gaze shifts for all participants also rules out a potential alternative explanation for the activation of the GFP during cube-rotation trials: face pareidolia. This is a perceptual effect where one sees a face in a pattern made from arbitrary elements that resembles the structural properties of a face^41,42^. In this sense participants may have started to perceive eyes in the cubes after being exposed to the experiment for a while. If face pareidolia was the cause, however, the activation associated with cube rotations would be expected to be smaller and more variable across participants^43^. In a recent fMRI study^44^ we had already demonstrated that a gaze cue which does not involve any apparent visual motion is not sufficient to reproduce the GFP. The present study provides an explanation for this finding.

Having learned that gaze shifts are not the only stimuli, GFP activity is positively correlated with, we investigated whether it can be functionally dissociated from brain areas specialized in the processing of visual motion, delineated by previous work resorting to a large spectrum of visual motion stimuli and usually summarized under terms like MT+ or the “visual motion complex”^45–48^ (see Figure 2). To this end, in Experiment 2 we first determined an MT+ ROI for each participant using a standard MT+ localization paradigm involving passive viewing of moving and static dots. We compared the responses to *cube-rotation* and *gaze-shift following* within and between the GFP and the MT+ ROIs. Importantly, in Experiment 2 we had modified the paradigm so that participants only saw either the gaze shift or the cube rotation in each trial, whereas in Experiment 1 they were presented together (see Figure 1). We introduced this modification to be able to explain a potential outcome of Experiment 1, namely that the GFP would be active during *cube-rotation* trials but with smaller *β*-values than during *gaze-shift* trials. This hypothetical result could have been explained by the observation that behavioral responses to observed gaze shifts are reflexive and occur even if the task requires to ignore them^14,49,17,19^. By separating the stimuli, we therefore ensured that any BOLD response we would observe is a genuine response to the stimulus in question. However, because our final results of Experiment 1 yielded larger *β*-values related to *cube-rotation*, falsifying the hypothesis that the GFP is a cortical module dedicated to the processing of others’ gaze-direction, we did not continue with data collection until all quantitative analyses of experiment 2 reached the set threshold (see *Transparent Changes*). To analyze Experiment 2, we estimated the hemodynamic response functions (HRF) for the different trial types (grouped by stimulus and target ID) and ROIs and computed the correlation between them (see Table 3 for a list of all pairings and Figure 7 for a conceptual illustration). This method protected the analysis from effects that might arise from differences in the visual complexity of the stimuli, as it relies purely on temporal dynamics. A property of the experimental design led us to predict a specific response pattern that would occur if the GFP corresponded to area MT: As illustrated in Figure 7 during each trial of our experiments two motion events occurred, first the gaze shift/ cube rotation towards the target and second the reset to the neutral orientations at the end of each trial. Importantly, the second motion event happened during the period in which the participants looked at the respective target object, meaning that it occurred in their visual periphery. As is well-known, the areas of the motion complex show lateralized responses to peripheral input. Therefore, we predicted that if ROIs consist of voxels which are part of MT+, they should exhibit sustained responses in the contralateral as compared to the ipsilateral hemisphere relative to the second motion event (see Table S1 and^37^). This would give rise to a specific correlation pattern among the pairs (yellow and blue groups in Figure 5 and Table 3). However, our data do not provide sufficient evidence to confirm or refute this prediction as we terminated data collection before having reached the set threshold, as explained above. Instead, we found *strong evidence* in favor of the alternative prediction that the correlations depend on the condition. That is, correlations between BOLD responses during trials of the same condition were slightly stronger than between those of different conditions (black and red groups in Figure 5 and Table 3). However, it is essential that this was the case for both ROIs and not just for the GFP, as the property of domain-specificity would require. The fact that both ROIs showed this feature suggests, firstly, that GFP and MT+ cannot be distinguished on the basis of their response properties in our experiment, and secondly, that the two stimuli induce similar – the cross-condition correlations are well above 0.5 on average (Figure 5, red bars) – yet, distinct HRFs within a brain area known to process also low-level visual motion.

**Figure 7.**
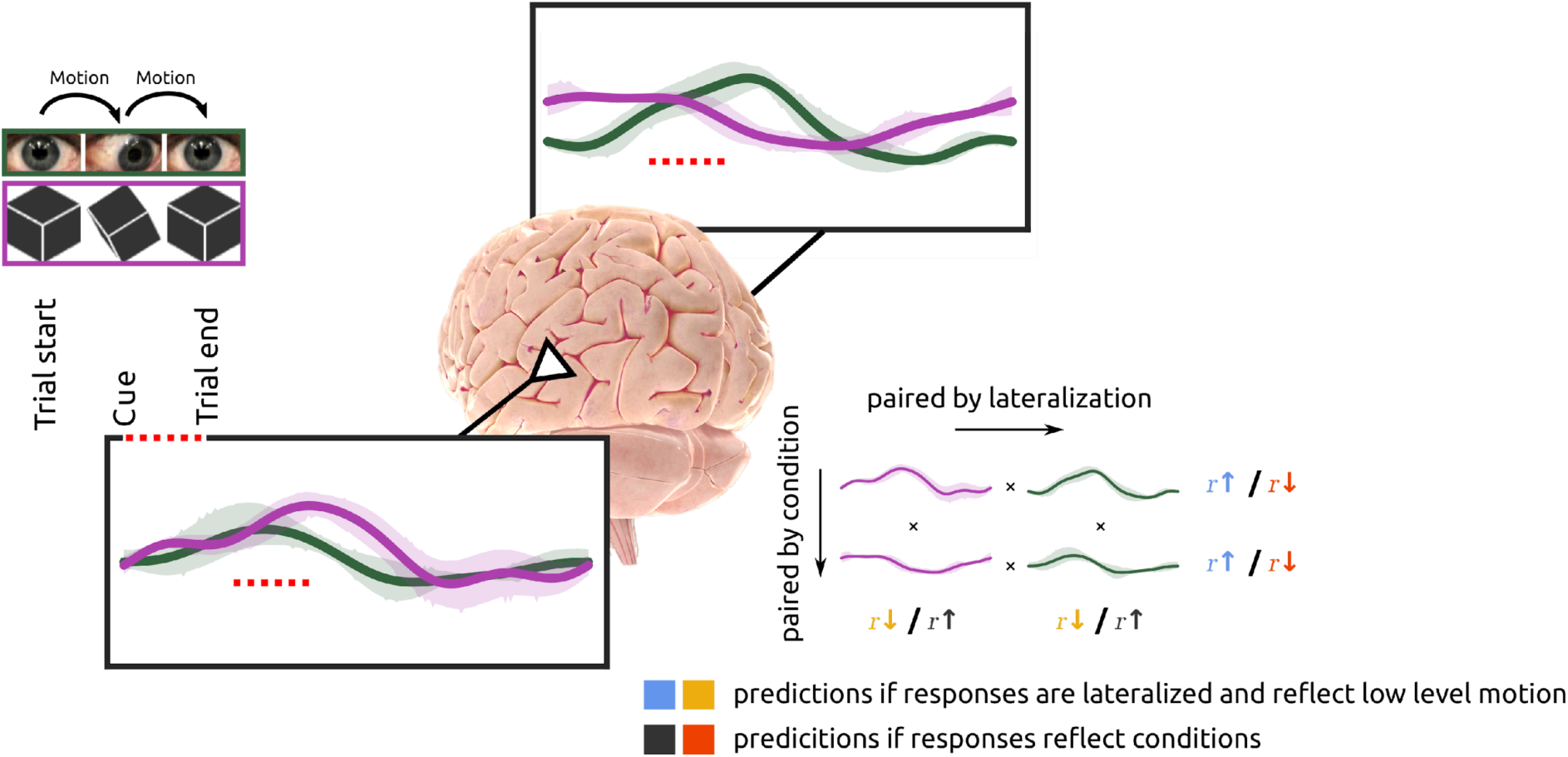
Illustration for Q3 and Q4. Shown are 4 example HRFs from one pilot subject to illustrate the analysis and predictions of experiment 2. In the illustrated gaze-only trial the participant looked at the right target such that the second motion event occurred in her left visual hemifield triggering a larger and longer lasting response in the right hemisphere compared to the left hemisphere (green curves). In the illustrated cube-only trial the participant fixated the left target leading to the opposite pattern (purple curves). Computing the correlations between the responses yields results that are in line with the blue and yellow predictions, indicating that the low-level motion components in the stimuli trigger lateralized responses. The shaded areas represent the 95% CI of the HRF estimates. See Table 3 for a full list of the predictions for all relevant response pairs.

Why did our data from the MT+ ROIs not match the pattern that we predicted based on properties of the motion complex regarding lateralization? We think the reason is that areas MST and FST and others that join area MT in the motion complex may comprise neurons with large receptive fields with no or only weak lateralization^47^. This is why they can be expected to exhibit activity evoked by both the central and the peripheral motion stimuli without much difference between hemispheres. Hence their contribution to the BOLD signals in the two ROIs may most probably have covered the much more specific contribution of area MT. Yet, irrespective of our inability to emphasize area MT, the similarity of the response patterns for both ROIs supports the localization of the GFP within MT+. This is also indicated by the strong correlation between the two ROIs, i.e., testing whether an HRF pair correlated (or not) in one ROI was also correlated (or not) in the other ROI (see Figure 5, scatter plot; Table S1, Q5).

Interestingly, the exploratory group level fits of the HRFs shown in Figure 4 reveal the pattern that was predicted based on lateralized responses independent of the experimental condition. Especially in the left hemisphere, BOLD responses during trials in which the target was on the right side were significantly reduced as compared to responses during trials in which the target was in the middle or on the left side (non-overlapping *CI*^95%^). Importantly, this pattern was present in both ROIs (top and bottom row in Figure 4) and for both experimental conditions (green and purple curves in Figure 4). In the right hemisphere the differences were less pronounced (*CI*^95%^do overlap), but met the prediction in that the opposing trend to that observed in the left hemisphere was present. Furthermore, when comparing the group level HRF estimates across ROIs and experimental conditions qualitatively, the similarity between all HRFs is striking.

Particularly interesting are the present results also in the light of studies which have suggested that biological motion is represented as a special category in parts of the temporal cortex^50–53^. While others have already suggested that these findings may merely represent quantitative rather than qualitative effects^54^ our results provide further evidence that it is rather task difficulty and context that determines the extent of activation of visual motion processing areas and not categorical membership. We base this argument on the fact that the GFP ROI that was defined through biologically meaningful motion, i.e. eye-movements, nevertheless exhibited larger *β*-values in relation to cube-rotations in experiment 1, but not in experiment 2 in which the stimuli were presented separately (see smaller cube-rotation related amplitudes in Figure 4). We think that two aspects are capable of explaining this seemingly contradictory finding, namely visual complexity and the ecological relevance of faces. Both may create a situation in experiment 1 in which the cube rotation is harder to detect (visual complexity of the face in the background) or utilize (attentional disengagement from the ecologically more relevant face), thus requiring a boost of the perceptual signal related to the rotating cubes relative to the gaze-cue. In contrast, Experiment 2 did not necessitate such a boost, as there was no competition between the stimuli.

If it is not the posterior temporal cortex that is tuned to socially guided spatial attention, are there other areas of the brain that are? In an exploratory analysis of experiment 1 we addressed this question by analyzing differences in the global brain activity associated with the *gaze-shift* and the *cube-rotation* conditions. To this end, we screened the *β*-maps for voxels exclusively associated with gaze-shifts and cube-rotations, respectively. Exclusively means that all voxels displayed in green in Figure 6 were positively correlated with the *gaze-shift* condition on the one hand, and negatively or not significantly correlated with the *cube-rotation* condition on the other (vice versa for the purple shaded voxels). In this comparison activity associated with the gaze-shifts was predominantly located in early visual areas, likely due to the higher visual complexity of the behaviorally relevant stimulus. Some smaller hotspots were in the temporal cortex (BA 21, 22 and 41) which are functionally associated with auditory processing and speech perception (based on neurosynth^36^ associations), possibly reflecting the need to process the verbal instructions. But why should there be larger activity evoked by the instruction to exploit gaze? The reason might be that we as social beings are well trained to use other peoples’ gaze. Hence, the activations associated with the instructions may not be suppressed by the less demanding gaze cues, yet by the cube cues whose geometrical interpretation may be more challenging. Figure S2 and S3 provide evidence supporting this explanation, as it shows that in most participants the *cubes-rotation* related *β*-values are negative for the two example ROIs considered. The most remarkable distinction of the activity exclusively related to the *cube-rotation* condition is the prominent activation of parietal areas (BA 39, 40, 7). This is surprising insofar as the hallmark of gaze-following is the redirection of visuospatial attention to specific spatial locations, a behavior known to involve the dorsal attention network (DAN)^55–59^. However, here, our results revealed that the IPS, a central node of the DAN, is not activated in the *gaze-following* condition, but in the *cube-rotation* condition, which is identical in behavioral and geometrical terms, the very area is active (see also Figure S4). Thus, it seems that gaze-following behavior as required in this experiment does not depend on the dorsal attention network which is at odds with previous findings^60,25,21^. One potential explanation for the absence of parietal activity might be that while extracting directional information from the cube-rotation requires some form of mental rotation, gaze-shift following does not. Mental rotation and viewpoint-dependent object recognition has been reported to involve parietal activity^61,62^. However, this does not explain why parietal activity has been observed in relation to gaze-following in previous studies^28,63^. A difference between the current paradigm and previous versions may offer an insight. Namely, while in previous studies participants had to delay their response to the cues until a dedicated go-signal appeared after a fixed time, in the present version participants were asked to respond to the cue immediately. Thus, previous designs^28,31^ offered a constant period between the cue and the go-signal in which the planning of the response saccade unfolded which might have triggered a noticeable modulation of the BOLD signal in the posterior parietal cortex^64,65^. Thus, previously observed gaze-following related parietal activity may have been related to intentional planning, an aspect not present in the present experimental design.

In conclusion the findings of the present study force us to abandon previous hypotheses about the modular processing and the categorical specificity of the gaze of others as a spatial reference^1,3,8^. Rather, we suggest that on the perceptual level core pathways, for example those which process visual motion, are exploited. Importantly, these conclusions have to be seen in the context of previous studies that discovered and investigated the GFP^8,28,29, 30,31,32,33,34,38^. Because of the small sample size, the exploratory analysis of the present study is limited and only serves to illustrate global activity patterns. However, any other specific brain area that might be an EDD, aside from the GFP, would have been discovered in any of the previous studies which were designed to capture the full behavioral relevance of active gaze following. Since the GFP was the only direct candidate neural substrate for the EDD, and we show here that its response is not specific to gaze stimuli, the broader conclusion that a domain-specific EDD does not exist is justified. This result is in line with recent findings which suggest that other perceptual categories, like faces^66,67^ or actions ^68^ have no modular or domain-specific representation, as well. At other levels of processing, there again appear to be differences between the processing of others’ gaze and geometric objects in terms of spatial cues, as indicated by the lack of parietal activity in the case of the former. Whether the reason for this is a more fundamental difference between these two types of cues (see the ‘association hypothesis’ suggested by Vecera and Rizzo^69^) or merely a consequence of overtraining to exploit the gaze of others cannot be answered here.

## Materials and Methods

### Transparent changes

Because experiment 1 already provided clear evidence regarding the main question of this study (Q6a, see Table S1) we stopped data collection before reaching the evidence threshold set in the registered research plan^37^ for Q3.1 & 4.1.

We added group-level fits of the HRFs for experiment 2 not described in the research plan^37^ (Figure 4).

The data of participant 6 was not included in the analysis regarding questions 3, 4 and 5 (see Table S1) because for this participant no individual GFP ROI could be delineated. Including this participant’s data, however, only had a quantitative effect on the results.

In the research plan^37^ we stated that results regarding Q3.1&2 that deviate from the predictions of lateralized responses would invalidate the analysis of experiment 2. This statement was based on the assumption that the motion localizer paradigm would delineate only those areas of the MT+ complex that are strictly lateralized. However, this assumption was not justified because we did not limit the low-level motion stimulus to contralateral stimulation but stimulated the whole visual field. Thus, the delineated ROI arguably comprises neighboring motion sensitive areas like MST and FST with no or the opposite lateralization. In the present manuscript we deleted this restriction, changed Table S1 accordingly and provide an interpretation of the results.

### Ethical Approval

The research presented here was approved by the Ethics Board of the Medical Faculty, University of Tübingen, Germany and was conducted in accordance with the principles of human research ethics of the Declaration of Helsinki. Written consent was obtained from all participants, and they were compensated with a lump sum of 40€.

### Sampling plan

We employed Bayesian methods for analysis. In line with suggested guidelines for registered reports (e.g.^70^) we included further participants the relevant Bayes Factors reached the threshold for *strong evidence* (≥ 10) in favor of the null-or the alternative hypothesis. Note that throughout this manuscript Bayes Factors are reported as *log*_10_ transforms, although this is not specified everywhere to improve readability. Furthermore, Bayes Factors are always reported from the perspective of the null hypothesis (*BF*_01_), this implies that values above 1 correspond to *strong evidence* in favor of the null hypothesis and values below −1 in favor of the alternative hypothesis. The exclusion of data was based solely on the observed behavior of the subject during the experiment. We observed the participants performing the tasks with a camera and recorded their answers, i.e., the direction of their response saccades, in writing for each trial. This was necessary since eye-tracking quality within the fMRI scanner is often not stable. We sampled data from 7 right-handed participants, 4 females and 3 males with a mean age of 29.6 ± 11.2 years. In accordance with the registered sampling plan^37^, we excluded the data of two participants before conducting an analysis because they had incorrect responses of more than 10% in at least two runs. We excluded the data from one run of another participant because more than 10 % of the answers were incorrect.

### Setup and general procedure

The study consisted of two experimental and one localizer tasks distributed over two separate fMRI sessions on different days. We acquired MR images using a 3T scanner (Siemens Magnetom Prisma) with a 20-channel phased array head coil. The head of the subjects was fixed inside the head coil by using plastic foam cushions to avoid head movements. An AutoAlign sequence was used to standardize the alignment of images across sessions and subjects. A high-resolution T1-weighted anatomic scan (MP-RAGE, 176 × 256 × 256 voxel, voxel size 1 × 1×1 mm) and local field maps were acquired. Functional scans were conducted using a T2*-weighted echo-planar multibanded 2D sequence (multiband factor = 2, TE = 35 ms, TR = 1500 ms, flip angle = 70°) which covered the whole brain (44 × 64 × 64 voxel, voxel size 3 × 3 × 3 mm, interleaved slice acquisition, no gap).

Participants were positioned in the scanner and viewed the stimulus monitor that was placed at its head-end via a mirror system. The distance between the participant and the monitor was ∼190 cm and it covered ∼20° of the field of view in the horizontal and ∼12° in the vertical. In the session with experiment 1 participants wore air pressure headphones. The stimuli and data collection was controlled via a custom software package^71^.

### Experiment 1

In each trial of Experiment 1 (see Figure 1) participants saw a photograph of the face of a female with superimposed cubes and a red fixation dot positioned on the nasion. The size of the cubes matched the size of the irises, and they were arranged so that they are equidistant from the nasion in the vertical direction as the eyes are in the horizontal direction. Each trial began with a verbal instruction delivered over headphones during which the face looked towards the participant, her iris color was grayish, and the cubes pointed straight ahead with one of their vertices. The verbal instruction was either *Gaze*, *Cubes,* or *Iris* indicating which of the upcoming cues had to be used to identify the target rectangle among three differently colored rectangles at the bottom of the stimulus image. After a variable ITI between 2 and 5 s, during which participants had to fixate the central fixation dot, the face made a gaze shift towards one of the rectangles, her iris color changed to match one of the target colors and the cubes rotated so that the frontal vertex pointed towards one of the targets. The participants had to respond to these cues as quickly as possible with a saccade to the target and maintain the fixation until the start of the next trial. The start of the next trial was indicated by resetting the cues to their initial state and by the next instruction. Each trial lasted between 5.5 and 8.5 s and participants completed 7 runs of 81 trials each. During each run, targets and instructions were randomized. Participants were allowed to rest between trials.

### Experiment 2

In Experiment 2, the face and the cubes were presented separately (*gaze-only*, *cubes-only*). Here, the *Iris-color* condition was omitted (see Figure 1). Therefore, the instruction at the beginning of each trial was not necessary and was omitted. In each run, only one type of cue (*Gaze* or *Cubes*) was presented, but the targets were randomized, and participants completed 4 runs of 81 trials each. Otherwise, the task and the temporal sequence were the same as in Experiment 1.

### Motion localizer

To localize the complex of motion processing areas (MT+) participants performed a standard motion localizer task in which they alternately viewed dot motion and a static dot pattern while maintaining fixation on a central fixation dot. Both the moving and the static dots were presented 15 times per run for 15 seconds each. In total, participants performed 6 runs. The dot field had a size of 7 deg and consisted of 200 dots with a dot size of 0.1 deg and a lifetime of 500 ms. The lifetimes of individual dots were asynchronous. During the motion period, individual dots moved at a speed of 1.5 deg/s. In 3 out of the 6 runs the movement direction of individual dots was randomized (noise level of 100%) and in the other 3 runs, the dots moved coherently (noise level of 0%) in a direction randomly drawn without replacement from a vector specifying 45 possible directions, uniformly sampling 360 deg.

### Registered Analysis

Table S1 provides an overview of all preregistered questions and related hypotheses, while a detailed description can be found in reference^37^. Some changes were necessary and are reported in the *Transparent Changes* section.

### Preprocessing

We used the software package *fMRIPrep* (ver. 20.2.1)^72,73^ (RRID:SCR_016216) which is based on *Nipype* 1.5.1^74^ (RRID:SCR_002502) to preprocess the fMRI data. The following sections describing the preprocessing are boilerplates automatically generated by *fMRIPrep* and inserted unchanged.

### Anatomical data preprocessing

A total of 1 T1-weighted (T1w) images were found within the input BIDS dataset. The T1w image was corrected for intensity non-uniformity (INU) with N4BiasFieldCorrection^75^, distributed with ANTs 2.3.3^76^ (RRID:SCR_004757), and used as T1w-reference throughout the workflow. The T1w-reference was then skull-stripped with a *Nipype* implementation of the antsBrainExtraction.sh workflow (from ANTs), using OASIS30ANTs as target template. Brain tissue segmentation of cerebrospinal fluid (CSF), white-matter (WM) and gray-matter (GM) was performed on the brain-extracted T1w using fast^77^ (FSL 5.0.9, RRID:SCR_002823). Brain surfaces were reconstructed using recon-all^78^ (FreeSurfer 6.0.1, RRID:SCR_001847), and the brain mask estimated previously was refined with a custom variation of the method to reconcile ANTs-derived and FreeSurfer-derived segmentations of the cortical gray-matter of Mindboggle^79^ (RRID:SCR_002438). Volume-based spatial normalization to one standard space (MNI152NLin2009cAsym) was performed through nonlinear registration with antsRegistration (ANTs 2.3.3), using brain-extracted versions of both T1w reference and the T1w template. The following template was selected for spatial normalization: *ICBM 152 Nonlinear Asymmetrical template version 2009c*^80^ (RRID:SCR_008796; TemplateFlow ID: MNI152NLin2009cAsym).

### Functional data preprocessing

For each of the runs found per subject (across all tasks and sessions), the following preprocessing will be performed. First, a reference volume and its skull-stripped version were generated by aligning and averaging 1 single-band references (SBRefs). A B0-nonuniformity map (or *fieldmap*) was estimated based on a phase-difference map calculated with a dual-echo GRE (gradient-recall echo) sequence, processed with a custom workflow of *SDCFlows* inspired by the epidewarp.fsl script^81^ and further improvements in HCP Pipelines^82^. The *fieldmap* was then co-registered to the target EPI (echo-planar imaging) reference run and converted to a displacements field map (amenable to registration tools such as ANTs) with FSL’s fugue and other *SDCflows* tools. Based on the estimated susceptibility distortion, a corrected EPI (echo-planar imaging) reference was calculated for a more accurate co-registration with the anatomical reference. The BOLD reference was then co-registered to the T1w reference using bbregister (FreeSurfer) which implements boundary-based registration^83^. Co-registration was configured with six degrees of freedom. Head-motion parameters with respect to the BOLD reference (transformation matrices, and six corresponding rotation and translation parameters) are estimated before any spatiotemporal filtering using mcflirt^84^ (FSL 5.0.9). BOLD runs were slice-time corrected using 3dTshift from AFNI 20160207^85^ (RRID:SCR_005927). First, a reference volume and its skull-stripped version were generated using a custom methodology of *fMRIPrep*. The BOLD time-series (including slice-timing correction when applied) were resampled onto their original, native space by applying a single, composite transform to correct for head-motion and susceptibility distortions. These resampled BOLD time-series will be referred to as *preprocessed BOLD in original space*, or just *preprocessed BOLD*. The BOLD time-series were resampled into standard space, generating a *preprocessed BOLD run in MNI152NLin2009cAsym space*. Several confounding time-series were calculated based on the *preprocessed BOLD*: framewise displacement (FD), DVARS and three region-wise global signals. FD was computed using two formulations following Power (absolute sum of relative motions^86^) and Jenkinson (relative root mean square displacement between affines^84^). FD and DVARS are calculated for each functional run, both using their implementations in *Nipype* (following the definitions by Power^86^). The three global signals are extracted within the CSF, the WM, and the whole-brain masks. Additionally, a set of physiological regressors were extracted to allow for component-based noise correction (*CompCor*^87^). Principal components are estimated after high-pass filtering the *preprocessed BOLD* time-series (using a discrete cosine filter with 128s cut-off) for the two *CompCor* variants: temporal (tCompCor) and anatomical (aCompCor). tCompCor components are then calculated from the top 2% variable voxels within the brain mask. For aCompCor, three probabilistic masks (CSF, WM and combined CSF+WM) are generated in anatomical space. The implementation differs from that of Behzadi et al.^87^ in that instead of eroding the masks by 2 pixels on BOLD space, the aCompCor masks are subtracted a mask of pixels that likely contain a volume fraction of GM. This mask is obtained by dilating a GM mask extracted from the FreeSurfer’s *aseg* segmentation, and it ensures components are not extracted from voxels containing a minimal fraction of GM. Finally, these masks are resampled into BOLD space and binarized by thresholding at 0.99 (as in the original implementation). Components are also calculated separately within the WM and CSF masks. For each CompCor decomposition, the *k* components with the largest singular values are retained, such that the retained components’ time series are sufficient to explain 50 percent of variance across the nuisance mask (CSF, WM, combined, or temporal). The remaining components are dropped from consideration. The head-motion estimates calculated in the correction step were also placed within the corresponding confounds file. The confound time series derived from head motion estimates and global signals were expanded with the inclusion of temporal derivatives and quadratic terms for each^88^. Frames that exceeded a threshold of 0.5 mm FD or 1.5 standardised DVARS were annotated as motion outliers. All resamplings can be performed with *a single interpolation step* by composing all the pertinent transformations (i.e., head-motion transform matrices, susceptibility distortion correction when available, and co-registrations to anatomical and output spaces). Gridded (volumetric) resamplings were performed using antsApplyTransforms (ANTs), configured with Lanczos interpolation to minimize the smoothing effects of other kernels^89^. Non-gridded (surface) resamplings were performed using mri_vol2surf (FreeSurfer).

### Copyright Waiver

The above boilerplate text was automatically generated by *fMRIPrep* with the express intention that users should copy and paste this text into their manuscripts *unchanged*. It is released under the CC0 license.

### Run- and first-level GLMs, ROI definition

For each participant we computed GLMs (general linear models) for each run individually (run-level) as well as a GLM for the combination of all runs (first-level) to obtain the respective *β*-images for each condition using the Python package *nilearn*^90^. To mitigate the effects of motion artifacts and other noise sources, the nuisance regressors *global_signal*, *csf*, white_matter, *trans_x*, *trans_y*, *trans_z*, *rot_x*, *rot_y*, *rot_z* and their respective first derivatives estimated by *fMRIPrep* were included in the design matrices. As a model for the *HRF* we used the *glover + derivative + dispersion* model provided by *nilearn* and included a polynomial drift model of order 3 to remove slow drifts in the data. Further, we masked the data with each run’s mask image (or the average for the first-level GLM) provided by *fMRIPrep* and applied smoothing with *fwhm* = 5 mm.

To define the GFP and the MT+ ROI the resulting first-level *β*-images were used to compute the contrasts *gaze-shift* minus *iris-color* (experiment 1) and *motion* minus *static* for the localizer task, respectively. For each participant these ROIs were determined individually by applying the most liberal statistical threshold that allowed to differentiate a separable activity patch at the location of the reference coordinates. These are for the GFP (*x* = −55, *y* = −68, *z* = 1) and (*x* = 50, *y* = −64, *z* = −3) ^31^ and for MT+ (*x* = −45, *y* = 70, *z* = 3) and (*x* = 46, *y* = −67, *z* = 4). The latter coordinates are the center of the output of a neurosynth.org database ^36^ search for *visual motion*. All coordinates reported in this study refer to the MNI space. The GFP-ROI was not identifiable for one participant in the way described above. For this one we constructed a spherical ROI with a radius of 8 mm located at the individual local maximum closest to the reference coordinates.

### Question 1/ 2: Comparison of gaze-shift and cube-rotation related activity in the left and right GFP

We extracted each participant’s run-level *gaze-shift* and *cube-rotation* related *β*-values from the GFP and averaged across voxels. Thus, we obtained 6 values per hemisphere and participant for each condition. These were fed into a Bayesian hierarchical regression model with *run* as group-level intercepts and slopes, and condition (*Question 1*) or hemisphere (*Question 2*) as the population-level effect. To avoid model fitting problems, *participant* was not added as a group-level effect becausethe number of participants was 5 as it is recommended by different sources (e.g. Bolker et al.^91^). Modeling was carried out in *R*^92^ using the *brms*^93^ package. Priors for the generative model were tuned to yield reasonable prior and posterior predictions for the pilot data (reported in reference^37^). As marginal prior for the effect of condition/ hemisphere on the *β*-values we used normal distribution with *μ* = 0, *σ* = 1. The *R* code of the model and the priors are reported in the Supplementary Materials section. The model provided an effect estimate, denoted as 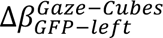. A positive value indicates that the gaze-shift related activity is larger than the cube-rotation related activity, while a negative value indicates the opposite. Bayes Factors were computed via the Savage-Dickey density ratio evaluating the ration between the posterior and the prior at the parameter value of zero^94^ and *log*_10_transformed. The terminology used in this study to describe the amount of evidence is adopted from Wetzels et al^95^.

### Question 3-5: HRF estimation and correlation analysis

For each participant and ROI, we estimated the *HRF* for each condition and target of experiment 2. This yielded 24 HRFs estimates in total for each participant (2 ROIs × 2 hemispheres × 2 conditions × 3 targets). To do so, the mean BOLD signals from the ROIs was extracted and linear deconvolution was applied to disentangle the responses to the different conditions/ targets. Prior to signal extractions, the BOLD images were denoised using the same confounds as for the GLMs (see above), smoothed with *fwhm* = 5 mm, and standardized. For the deconvolution, we resorted to the *nideconv* package^96^ and used 9 *fourier* regressors as the basis set which allows a larger flexibility than the canonical HRF while being less complex than a FIR model.

For each ROI, we computed linear correlation coefficients (employing the function *correlationBF*^97^) between all relevant pairs (see Table 3) of the responses to the left, middle and right target within and across the conditions of experiment 2. In total, we obtained the correlation coefficients of 34 HRF-pairs for each participant and ROI. These were grouped by the expected correlation if the responses to the second motion cue in the end of trials were lateralized (color code of the circles in Table 3) and by within- and cross-condition pairings (color code of the dots in Table 3). Figure 7 shows an example. These groups were analyzed for differences using Bayesian hierarchical models. The correlation coefficients were modeled as gaussian distributions within the limit of possible values of correlation coefficients – between −1 and 1. Priors for the generative model were tuned to yield reasonable prior and posterior predictions for the pilot data (see reference^37^). As marginal priors for the effect of the respective groupings we used a normal distribution with *μ* = 0 and *σ* = 2. The first model provided the effect estimate as 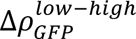 which is negative if the prediction based on lateralization is met. The second model will provide the effect estimate as 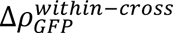 which is positive if the prediction based on condition is met. Bayes Factors were computed via the Savage-Dickey method and *log*_10_transformed.

Finally, if the correlation of a pair of HRFs is high in one ROI, we tested whether it is then also high in the other ROI (Q5). We did so by computing the correlation between the ROIs and the Bayes Factor testing against no correlation using the *correlationBF* function.

### Exploratory Analysis

To explore brain wide differences between *gaze-shift* and *cubes-rotation* following related activity we computed second level ***β***-maps for the two conditions based on the non-smoothed first level ***β***-maps from experiment 1. The resulting second level map was smoothed using a 8 mm kernel. To assess qualitative as well as quantitative differences between activity related to the conditions in a first step using the thresholded ***β***-maps (*p* ≤ 0.05, uncorrected, minimum cluster size of 15 voxels) we created masks that excluded all significantly positively correlated voxels. In the second step we applied to the thresholded ***β***-maps of respective other condition. This means we removed all voxels that are positively correlated with the *cube-rotation* condition from the ***β***-map of the *gaze-shift* condition and vice versa. We applied a cluster threshold of 15 voxels to the results once more to partially account for the very liberal alpha threshold used here. This allowed us to create ***β***-maps that contain only genuinely activated areas for the two conditions (Figure 6).

## Acknowledgements

We thank Dr. Friedemann Bunjes for his invaluable technical assistance.

We acknowledge support from the German Research Foundation (DFG) project TH425/17-1 and from the Open Access Publication Fund of the University of Tübingen.

## Author contribution

MG designed the study, collected the data, analyzed the data, wrote the manuscript; PD collected the data; PT designed the study, wrote the manuscript

## Competing interests

The authors declare no competing interests.

## Data Availability Plan

Unprocessed data organized in accordance with the BIDS format is publicly available at doi:10.18112/openneuro.ds005903.v1.0.1.

Any code used for analysis is publicly available at https://github.com/maalaria/fMRIus/tree/gaze-motion.

## Notes

doi:10.18112/openneuro.ds005903.v1.0.1

### Competing Interest Statement

The authors have declared no competing interest.

### Summary of Updates

Minor revision of the the Abstract and Discussion

