## Supplementary Material for "Evidence against a cortical module for processing communicative gaze"

**Table S1.** Specification of preregistered hypotheses

| Question | Null Hypothesis | Outcome Measures | Sampling plan | Analysis Plan | Interpretation of outcomes |
|---|---|---|---|---|---|
| **Q1** Is the GFP-ROI differentially activated during *Gaze* and *Cubes* in experiment 1? | $H_0^{Q1.1}: \Delta\beta_{GFP-left}^{Gaze-Cubes} = 0$<br>$H_0^{Q1.2}: \Delta\beta_{GFP-right}^{Gaze-Cubes} = 0$ | Beta estimates of BOLD signal | $BF_{01/10}^{Q1} \geq 10$ | Bayesian hierarchical model | **1.** $\Delta\beta_{GFP}^{Gaze-Cubes} \leq 0$: no difference between *Gaze* and *Cubes* related activity or less *Gaze* related activity in the GFP. Support against the hypothesis that the GFP is a specialized module for gaze-following.<br>**2** $\Delta\beta_{GFP}^{Gaze-Cubes} > 0$: The GFP is stronger activated during *Gaze* trials than during *Cubes* trials. Either the GFP is a specialized gaze-following module or it is a subpart of MT+ reflecting retinotopic locations. These possibilities are disentangled in Q6. |
| **Q2** Is there a hemispheric dominance for *Gaze* or *Cubes*? | $H_0^{Q2.1}: \Delta\beta_{Gaze}^{RH-LH} = 0$<br>$H_0^{Q2.2}: \Delta\beta_{Cubes}^{RH-LH} = 0$ | Beta estimates of BOLD signal | $BF_{01/10}^{Q2} \geq 10$ | Bayesian hierarchical model | **1.** $\Delta\beta_{Gaze/Cubes}^{RH-LH} = 0$: no difference between the left and right hemisphere.<br>**2.** $\Delta\beta_{Gaze/Cubes}^{RH-LH} > 0$: The right hemisphere is stronger activated than the left hemisphere. Complies with the right hemispheric dominance of visual face processing.<br>**3.** $\Delta\beta_{Gaze/Cubes}^{RH-LH} < 0$: The left hemisphere is stronger activated than the right hemisphere. |

| | | | | | |
|---|---|---|---|---|---|
| **Q3.1** Does target ID in conjunction with hemisphere affect the level of correlations in MT+. | $H_0^{Q3.1}: \Delta\rho_{MT+}^{low-high} = 0$ | Correlation coefficients of HRF estimates of BOLD signals | $BF_{01/10}^{Q3.1} \geq 10$ | Bayesian hierarchical model | **1.** $\Delta\rho_{MT+}^{low-high} = 0$: no lateralized response in MT+. ROI does not only contain araes with contralateral receptive fields or the peripheral motion cue is not of relevance.<br>**2.** $\Delta\rho_{MT+}^{low-high} < 0$: the result matches the prediction based on lateralization.<br>**3.** $\Delta\rho_{MT+}^{low-high} > 0$: reverse effect as predicted (implausible). Invalidates analysis of experiment 2. |
| **Q3.2** Does condition affect the level of correlation in MT+. Q3.1 and 3.2 are positive controls for the second part of the analysis. | $H_0^{Q3.2}: \Delta\rho_{MT+}^{within-across} = 0$ | Correlation coefficients between HRF estimates of BOLD signals | $BF_{01/10}^{Q3.2} \geq 10$ | Bayesian hierarchical model | **1.** $\Delta\rho_{MT+}^{within-across} = 0$: correlations within conditions and across conditions are equally strong. Matches the prediction for a visual motion area.<br>**2.** $\Delta\rho_{MT+}^{within-across} < 0$: correlations within conditions smaller than across conditions (implausible).<br>**3.** $\Delta\rho_{MT+}^{within-across} > 0$: correlations within conditions larger than across conditions. Indicates a differential representation of the conditions within a domain-general brain area. |
| **Q4.1** Does target ID in conjunction with hemisphere affect the level of correlations in the GFP | $H_0^{Q4.1}: \Delta\rho_{GFP}^{low-high} = 0$ | Correlation coefficients between HRF estimates of BOLD signals | $BF_{01/10}^{Q4.1} \geq 10$ | Bayesian hierarchical model | **1.** $\Delta\rho_{GFP}^{low-high} = 0$: no lateralized response in GFP. Given that $\Delta\rho_{MT+}^{low-high} < 0$, we will conclude that the GFP is functionally different from MT+. See Q6 for full interpretation. |

| | | | | | |
|---|---|---|---|---|---|
| | | | | | **2.** $\Delta\rho_{GFP}^{low-high} < 0$: the result matches the prediction based on lateralization. Given that $\Delta\rho_{MT+}^{low} < 0$, we will conclude that the GFP and MT+ are functionally homologous. See Q6 for full interpretation.<br><br>**3.** $\Delta\rho_{GFP}^{low-high} > 0$: reverse effect as prediction (implausible). See Q6 for full interpretation. |
| **Q4.2** Does condition affect the level of correlation in the GFP | $H_0^{Q4.2}: \Delta\rho_{GFP}^{within-across} = 0$ | Correlation coefficients of HRF estimates of BOLD signals | $BF_{01/10}^{Q4.2} \geq 10$ | Bayesian hierarchical model | **1.** $\Delta\rho_{GFP}^{within-across} = 0$: both conditions elicit the same responses in GFP. Indicating that it is not a specialized module for the processing of gaze. See Q6 for full interpretation.<br><br>**2.** $\Delta\rho_{GFP}^{within-across} < 0$: lower correlation within- than across-conditions (implausible).<br><br>**3.** $\Delta\rho_{GFP}^{within-across} > 0$: across-conditions pairs are less correlated than within-condition pairs. Supporting that the GFP is a specialized module if $\Delta\rho_{MT+}^{within-across} \leq 0$. See Q6 for full interpretation. |
| **Q5** are the correlation coefficients from the GFP correlated with those from MT+ | $H_0^{Q4}: \rho_{MT+/GFP} = 0$ | Correlation coefficients | $BF_{01/10}^{Q5} \geq 10$ | Bayesian correlation test | **1.** $\rho_{MT+/GFP} = 0$: no correlation between MT+ and GFP. GFP is functionally segregated from MT+.<br><br>**2.** $\rho_{MT+/GFP} > 0$: GFP is (part of) MT+ |

| | | | | | |
|---|---|---|---|---|---|
| | | | | | **3.** $\rho_{MT+/GFP} < 0$: GFP and MT+ are anticorrelated (implausible). |
| **Q6** Is the GFP (a) a specialized module for the processing of gaze and (b) functionally segregated from MT+? | - | - | - | Combination of Q1,3,4,5 | In order to conclude that the GFP is a specialized module the following results must occur:<br>(a)<br>Q1: $\Delta\beta_{GFP}^{Gaze-Cubes} > 0$<br>(b)<br>Q3.1: $\Delta\rho_{MT+}^{low-high} < 0$<br>Q3.2: $\Delta\rho_{MT+}^{within-across} = 0$<br>Q4.1: $\Delta\rho_{GFP}^{low-high} = 0$<br>Q4.2: $\Delta\rho_{GFP}^{within-across} > 0$<br>Q5: $\rho_{MT+/GFP} = 0$ |
| **Q7** Is the GFP identical with or a part of MT+? | - | - | - | Combination of Q1,3,4,5,6 | In order to conclude that the relevant stimulus component triggering activity in the GFP is motion and that it cannot be distinguished from MT+ the following results must occur:<br>Q1: $\Delta\beta_{GFP}^{Gaze-Cubes} \leq 0$<br>Q3.1/ 4.1: |

| | | | | | |
|---|---|---|---|---|---|
| | | | | | $\Delta\rho_{MT+}^{low-high} < 0 > \Delta\rho_{GFP}^{low-high}$<br><br>Q3.2/ 4.2:<br><br>$\Delta\rho_{MT+}^{within-across} = 0 = \Delta\rho_{GFP}^{within-across}$<br><br>Q5: $\rho_{MT+/GFP} > 0$ |

3

### *Model specification*

**Table S2.** *R* code in *brms* syntax for the models and the priors.

<table>
<tr><th></th><th>Formula</th><th>Priors</th></tr>
<tr><td rowspan="5">Q1/2</td><td rowspan="5">$\beta$ ~ x + (x | run),<br>family=gaussian(link="identity"),<br>x = {condition, hemisphere}</td><td>normal(0, 5), class='Intercept'</td></tr>
<tr><td>normal(0, 1), class='b', coef={'conditionGaze', hemisphereRH}</td></tr>
<tr><td>normal(.5, .5), class='sd', coef='Intercept', group='run'</td></tr>
<tr><td>normal(.3, .5), class='sd', coef='conditionGaze', group='run'</td></tr>
<tr><td>lognormal(.5, .5), class='sigma'</td></tr>
<tr><td rowspan="3">Q3/4</td><td rowspan="3">r | trunc(lb=-1, ub=1) ~ x,<br>family='gaussian',<br>x = {lateralization, condition}</td><td>normal(0, 1), class='Intercept'</td></tr>
<tr><td>normal(0, 2), class='b'</td></tr>
<tr><td>lognormal(0, .5), class='sigma'</td></tr>
</table>

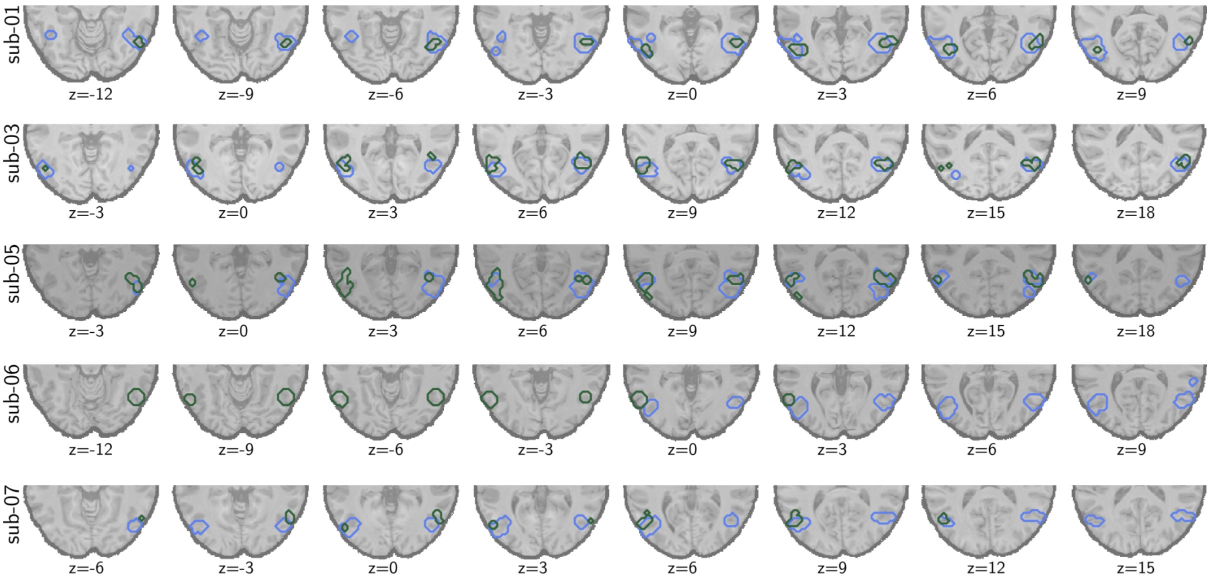


**Figure S1** Outlines of the GFP (green) and MT+ (blue) ROIs for each participant.

### *Language ROIs*

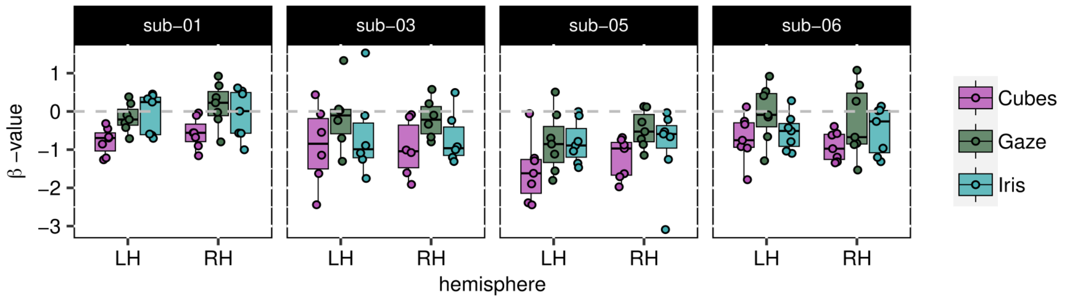


**Figure S2 Broca area.** $\beta$-values extracted from a ROI representing the Broca area. For all participants the values associated with the cube-rotation condition are smaller, and negative, than those associated with the gaze-shift condition. I.e. the activation of language areas does not reflect an activation of language areas as a simple BOLD contrast between the *gaze-shift* and *cube-rotation* conditions would suggest.

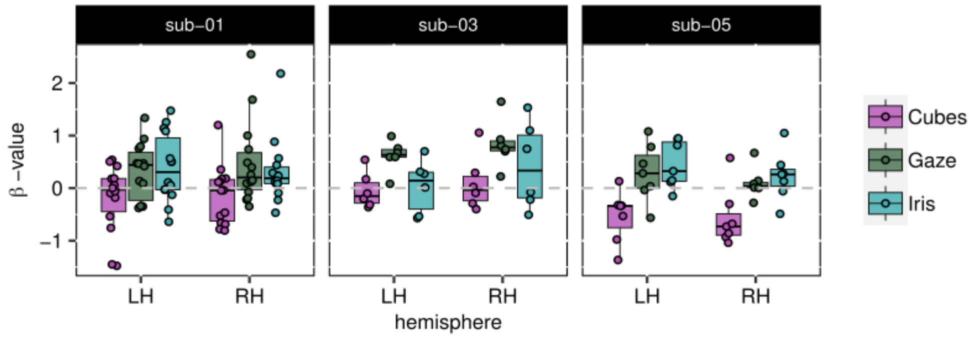


**Figure S3 Middle temporal gyrus.** Here the picture is less clear than for the Broca area. Subject 03 shows positive $\beta$-values associated with the *gaze-shift* condition while subject 05 shows negative $\beta$-values for the *cube-rotation* condition. The reason for this discrepancy might be the chosen ROIs are not actually representing comparable brain areas. For example it might be that the ROI used for subject 03 coincides with face-responsive areas in the temporal cortex while the ROI for subject 05 does not.

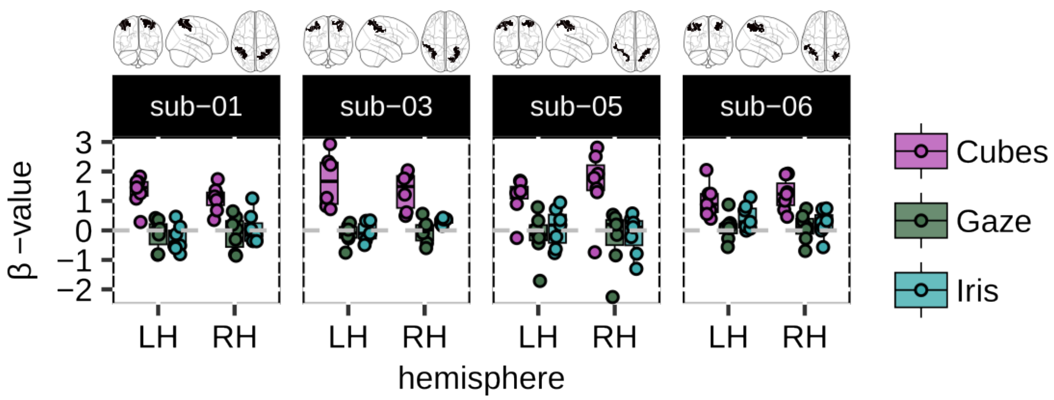

**Figure S4 Intraparietal sulcus** The ROIs were defined based on the contrast *gaze-shift following* minus *cube-rotation following*. The *β*-values related to the *gaze-shifts* (and *iris-color mapping*) are on average not different from 0.
